# Contaminant DNA in bacterial sequencing experiments is a major source of false genetic variability

**DOI:** 10.1101/403824

**Authors:** Galo A Goig, Silvia Blanco, Alberto L. Garcia-Basteiro, Iñaki Comas

**Affiliations:** Institute of Biomedicine of Valencia, IBV-CSIC, St. Jaume Roig 11, 46010, Valencia, Spain; Centro de Investigaçao em Saúde de Manhiça (CISM), Bairro Cambeve, Rua 12, Distrito da Manhiça, 1929 Maputo, Moçambique; Global Health Institute Barcelona, (ISGlobal), Rossello, 132, 4a, 08036 Barcelona, Spain; Amsterdam Institute for Global Health and Development, Amsterdam University Medical Centers, Paasheuvelweg 25, 1105 BP Amsterdam, Netherlands; CIBER in Epidemiology and Public Health, Madrid, Spain

## Abstract

Contaminant DNA is a well-known confounding factor in molecular biology and in genomic repositories. Strikingly, analysis workflows for whole-genome sequencing (WGS) data usually neglect the errors introduced by potential contaminations. We performed a comprehensive evaluation of the extent and impact of contaminant DNA in WGS by analyzing more than 4,000 bacterial samples from 20 different studies. We found that contaminations are pervasive and can introduce large biases in variant analysis. We showed that these biases can translate in hundreds of false positive and negative SNPs, even for samples with slight contaminations. Studies investigating complex biological traits from sequencing data can be completely biased if contaminations are neglected during the bioinformatic analysis. We used both real and simulated data to evaluate and implement reliable, contamination-aware analysis pipelines. Our results urge for the implementation of such pipelines as sequencing technologies consolidate as a precision tool in the research and clinical context.

## Introduction

Whole genome sequencing (WGS) has enhanced the study of complex biological phenomena in bacteria, such as population dynamics, host adaptation or outbreaks of microbial infections (*1, 2*). In addition, democratization of high-throughput sequencing technologies and continuous improvements in laboratory procedures are also turning WGS into a promising alternative for the clinical diagnosis and surveillance of several pathogenic species (*3*–*5*). Thus, many efforts in the basic and clinical research fields are directed to the improvement of bioinformatics pipelines to ensure the robustness of the conclusions drawn.

Central to many bacterial WGS bioinformatics pipelines is the identification of genetic variants. Incorrect identification of variants can have a major impact on several areas of microbiological research. Applications based on variant analysis include, but are not limited to, phylogenetics (*6*), phylodynamics and dating(*7*), genome-wide association studies(*8*), experimental evolution(*9*), epidemiological analyses(*10*) or drug development(*11*). Furthermore, the frequency at which each variant is observed in a sample can be used to characterize population genetics processes. Analysis of the allele frequency spectrum enables the study of population dynamics of clonal diversity within a niche or co-existence of mixed lineages(*12*). In the clinical field, variant analysis at a genomic scale allows the identification of pathogen species and genotypes, distinguish between relapse and superinfections, or prediction of resistance phenotypes and transmission links.

While many factors are taken into account when developing SNP calling pipelines, surprisingly the potential role of contaminants is seldomly considered (*13*). However, misinterpretation of contaminated data can lead to draw incorrect conclusions about biological phenomena (*14, 15*).

Genomic databases are known to encompass contaminated sequences, with assembled genomes that can contain large genomic regions from non-target organisms (*16, 17*). Strikingly, a recent study revealed that deposited bacterial and archaeal assemblies are contaminated by human sequences that created thousands of spurious proteins(*18*). While the potential impact of contaminants has been considered in fields like metagenomics or transcriptomics, most bacterial WGS analysis pipelines lack specific steps aimed to deal with contaminant data. This situation likely originates from the assumptions that microbiological cultures are mostly free of non-target organisms and that, if any, contaminating sequences hardly map to the reference genomes or are removed using standard filter cutoffs. To date, the extent of contaminations and their impact in bacterial re-sequencing pipelines has not been comprehensively assessed.

In this work, we use both real and simulated data to perform a detailed comparison of a standard bacterial mapping and SNP calling pipeline against two alternative contamination-aware approaches. First, we implement a taxonomic filter removing contaminant reads that allowed us to assess the extent of contaminations and estimate its impact in a dataset comprising 2,600 samples of 13 different species from 12 bacterial WGS projects. Second, we compare the performance of this taxonomic filter with a filter based on the similarity of the alignments, and evaluate the impact of contaminations in 8 WGS projects comprising 1,500 samples of *Mycobacterium tuberculosis* (MTB) WGS samples.

We found that contaminations are frequent across bacterial WGS studies and can introduce large biases in variant analysis despite using stringent mapping and variant calling cutoffs. Importantly, this is not only true for culture-free sequencing strategies, but also for experiments sequencing from pure cultures. We show that the effect size is not dependent on the amount of contamination and that samples with subtle contaminations can accumulate dozens of errors. We demonstrate that removing contaminant reads with a taxonomic classifier allows the implementation of highly accurate variant calling pipelines and provide a validated workflow for WGS analysis of MTB.

## Results

### Contaminations are common across WGS studies, even when sequencing from pure cultures

To assess the extent of contaminations across bacterial WGS studies, we taxonomically classified the sequencing reads of 4,194 WGS samples from 20 different studies using Kraken, a metagenomic read classifier that has been extensively used and evaluated in the literature. Out of these, 1,553 samples corresponded to *M. tuberculosis* sequencings, here referred as the *MTB dataset*, and 2,641 to other 13 bacterial species, here referred as the *bacterial dataset* (Table 1). According to taxonomic classifications, varying levels of contamination with non-target reads can be found in the different studies (Figure 1). From the *bacterial dataset, L. pneumophila, A. baumannii, L. monocytogenes, P. aeruginosa* and *N. gonorrhoeae* studies showed the expected taxonomic profile from pure culture isolate sequencings, since virtually all the reads are classified in their respective target genus. By contrast, contaminations can be clearly found in the rest of studies from this dataset, with an average of 45% of samples per study having less than the 90% of the reads coming from the target organism. The *T. pallidum* study represents an extreme case, with its samples having an average of only 40% of reads coming from this organism. This result is expected since in this study the samples were sequenced directly from clinical specimens using a bait capture strategy. However, high levels of contamination can be found in other studies where sequencing is performed from pure cultures (Figure 1a).

**Table 1.** Studies analyzed.

| Study name | Publication | Runs analyzed | Sample source | Dataset |
| --- | --- | --- | --- | --- |
| Mozambique | Unpublished | 138 | Clinical isolates | MTB<br>dataset |
| Kwazulu-Natal | Cohen et al.<br>2015 | 433 | Single colonies from clinical isolates | MTB<br>dataset |
| Nigeria | Senghore et al.<br>2017 | 73 | Clinical isolates | MTB<br>dataset |
| Belarus | Wollenberg et<br>al. 2017 | 552 | Clinical isolates | MTB<br>dataset |
| High-depth<br>sequencing | Trauner et al.<br>2017 | 63 | Clinical isolates | MTB<br>dataset |
| Sputum capture-sequencing | Brown et al. 2015 | 58 | Clinical respiratory specimens (culture-free sequencing with a bait capture strategy) | MTB dataset |
| Sputum direct-sequencing | Votintseva et al. 2017 | 68 | Clinical respiratory specimens (direct culture-free sequencing) | MTB dataset |
| MGIT sequencing* | Pankhurst et al. 2016 | 168 | Early-positive MGIT cultures (liquid) | MTB dataset |
| *A. baumannii* | Willems et al. 2016 | 36 | Single-colony recultured in broth | Bacterial dataset |
| *C. difficile* | Stone et al. 2016 | 54 | Pooled single-colony isolates | Bacterial dataset |
| Enterococcus† | Tyson et al 2018 | 197 | Isolates from retail meats | Bacterial dataset |
| *K. pneumoniae* | Holt et al 2015 | 285 | Human, animal and environmental isolates | Bacterial dataset |
| *L. monocytogenes* | Halbedel et al 2018 | 424 | Clinical isolates from human | Bacterial dataset |
| *L. pneumophila* | Timms et al 2018 | 48 | Pure culture isolates from human and cooling towers | Bacterial dataset |
| *N. gonorrhoeae* | Yahara et al 2018 | 272 | Pure culture isolates from human | Bacterial dataset |
| *P. aeruginosa* | Marvig et al 2015 | 445 | Clinical isolates from human | Bacterial dataset |
| *S. aureus* | Aanensen et al 2016 | 337 | Clinical isolates from 186 hospitals in 21 countries | Bacterial dataset |
| *S. enterica* | Gymoese et al 2017 | 366 | Human, animal and environmental isolates | Bacterial dataset |
| *T. pallidum* | Pinto et al 2016 | 25 | Clinical specimens (culture-free sequencing with a bait capture strategy) | Bacterial dataset |
| Vibrio‡ | Greig et al 2018 | 152 | Clinical isolates from human | Bacterial dataset |

\*This study included sequencings from non-MTB organisms. We analyzed the 168 reported as
MTB by the authors. †This study included sequencings from two species (*E. faecalis* and *E.* *faecium*). ‡This study included sequencings from different *Vibrio* species, we only analyzed the 152 reported as *V. cholerae* by the authors.

**Figure 1:**
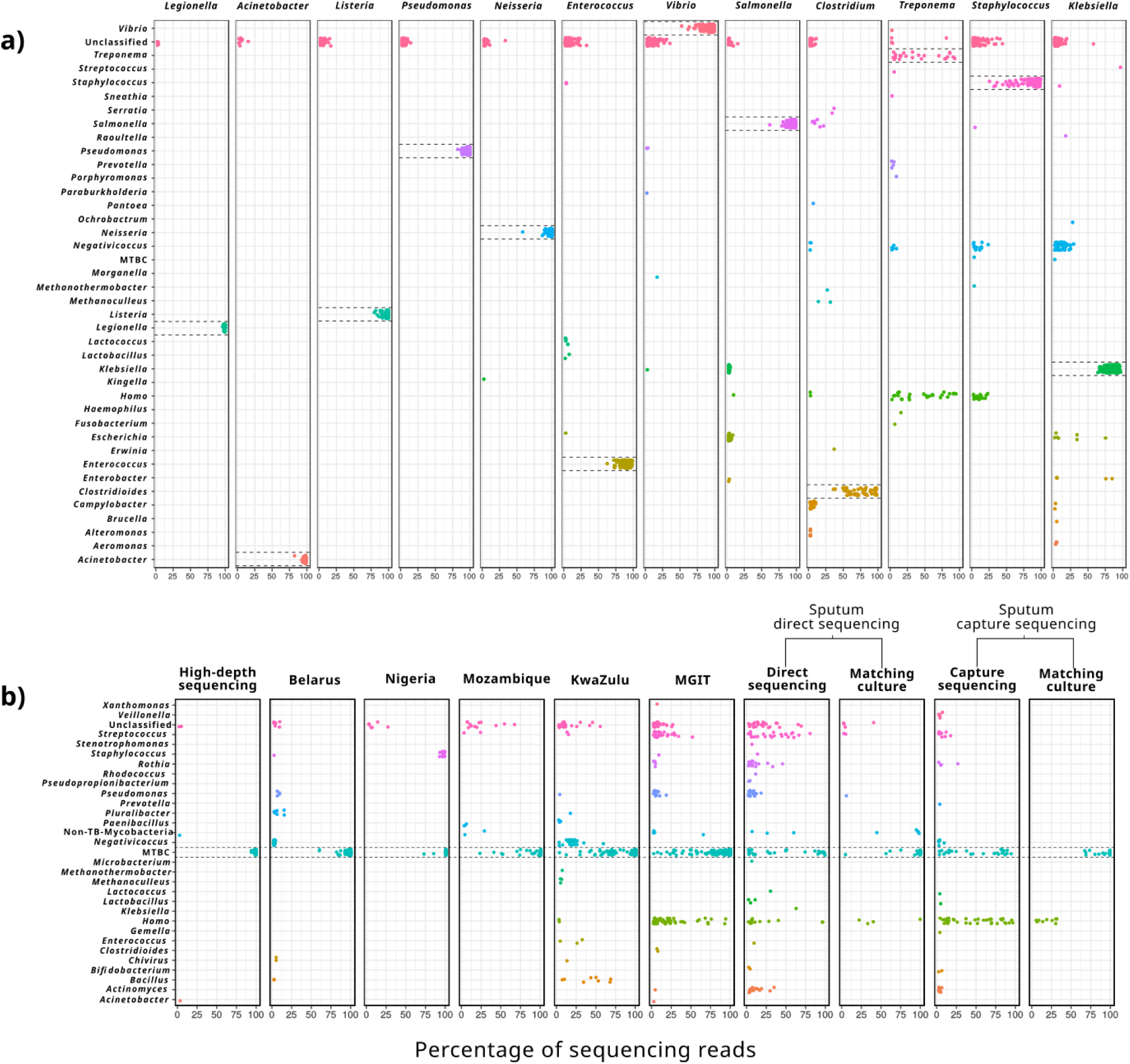
Proportion of sequencing reads for different organisms across 4,346 WGS samples from 20 different studies. Each dot represents a sample with a given percentage of sequencing reads coming from the genus indicated in the y-axis. Dashed lines highlight the target organism of each study. A 0.3 of vertical jitter was applied for better visualization. Only organisms in a proportion above the 2% are shown. a) Studies of the bacterial dataset. b) Studies of the MTB dataset. The two Enterococcus species analyzed in the bacterial dataset are shown under the same rectangle as they belong to the same genus and the same study.

When looking at the *MTB dataset*, we also observed contaminations to be common across studies (Figure 1b). As expected, direct sequencings from clinical specimens and early positive mycobacterial growth indicator tubes (MGIT), which are inoculated with primary clinical samples, present higher levels of contamination in terms of both the number of samples contaminated and the proportion of non-target reads within them. Common contaminants for these samples comprise human DNA, and bacteria usually found in oral and respiratory cavities like *Pseudomonas, Rothia, Streptococcus* or *Actinomyces*, and can constitute virtually all reads in some samples. However, as observed for the *bacterial dataset*, contamination was also detected in studies in which the sequenced DNA came from pure culture isolates. For instance, *Bacillus, Negativicoccus* and *Enterococcus* represented up to 68%, 58% and 32%, respectively, of different samples from the KwaZulu study. Strikingly, 17 out of 73 MTB samples from the Nigeria study were identified as *Staphylococcus aureus* (92% to 99% of reads), probably due to a mistake during data uploading or mislabeling in the laboratory. The high-depth dataset was mostly free of contamination, with the exception of two samples for which 3.32% of *Acinetobacter baumannii* and 2.83% of non-tuberculosis mycobacteria (NTM) was identified (representing 795,887 and 920,379 reads respectively).

### A taxonomic filter to selectively analyze non-contaminant reads

To assess the impact of these contaminations in bacterial WGS analysis, we compared the outcomes in variant calling for each sample before and after removing contaminant reads as classified by Kraken. We refer to this contamination removal methodology as “taxonomic filter” (detailed in Methods). To assess whether our Kraken setup can be safely used to remove contaminant reads across the analyzed datasets, we first estimated the proportion of reads that can be classified up to the level of species and genus for each organism using a simulated FASTQ file from the corresponding reference genome (Supplementary Table 1). For most of the organisms more than the 99% of the reads could be classified at species level for 250bp Illumina MiSeq sequencings (median = 99.35%) with the exceptions of *K. pneumoniae (*97.86%), *S. aureus* (95.01%) and *T. pallidum* (93.54%). For 100 bp Illumina HiSeq sequencings the proportion of reads classified for each organism was lower in every case (median=98.79%) with a dramatic drop for *T. pallidum* (72.74%), and with the exception of *M. tuberculosis* that remained 99.98%. At genus level, Kraken was able to classify most of the reads of each organism (median=99.89% for 250 bp sequencings; median=99.77% for 100 bp sequencings) with the exception of *S*.*aureus* that remained around 95% for both 250 and 100 bp sequencings. Interestingly, for *T. pallidum*, which showed to be the most difficult organism to classify at species level, 100% of reads were classified at genus level.

Second, we scanned all the WGS samples to estimate the maximum proportion of reads Kraken is capable of classify as the target organism in real samples (Supplementary Table 1). In most cases there was at least one sample per bacteria that could be classified as good as the reference genome (median difference between real and simulated sequencings of 1% at species level and 0.35% at genus level). The higher difference was observed for *T. pallidum* for which the maximum number of reads classified in a real WGS sample at genus level was of 94.75%.

Therefore, to safely analyze the effect that contaminants reads have in WGS sequencings of the *bacterial dataset*, we applied the taxonomic filter at the genus level, thus removing from each sample those reads classified as any other organism than the target genus (e.g we removed all non-*Acinetobacter* reads from the *A. baumannii* study). This strategy can safely remove contaminants at the cost of potentially analyzing reads from the same genus than the target organism. In addition, we avoided highly contaminated samples introducing extreme biases in the analysis by discarding samples with contaminations higher than 50% and depths lower than 40X (20X for *T. pallidum*, see Methods for a further explanation). From the initial 2,641 samples of the *bacterial dataset*, 2,233 met these criteria.

### Contamination impacts bacterial WGS analysis

The expected effect of contaminant read mappings is to produce mixed calls, leading to the identification of false positive variable SNPs (vSNPS). These false positive calls would alter the frequencies calculated at a given position, what might also produce false negative fixed SNPs (fSNPs) by lowering the frequency below the required cutoff to call fixed variants (90% frequency in this work). Overall, There was a high correlation between removing vSNPs and recovering fSNPs (Pearson Correlation Coefficient=0.76) (Figure 2). However, not all the contaminant reads are expected to affect positions with fSNPs and, in fact, for 405 samples (18%) the taxonomic filter removed the false positive vSNPs without affecting any fSNP. Similarly, in 38 samples (3%) we observed the recovery of at least one false negative fSNP without removal of vSNPs. Notably, we did not observe a correlation between the number of vSNPs removed and the degree of contamination of a sample (Pearson Correlation Coefficient = −0.06) (Table 2). This result suggests that the impact in variant analysis is highly dependent on which are both the contaminant and the target organisms, rather than the amount of contaminating reads.

**Table 2.**
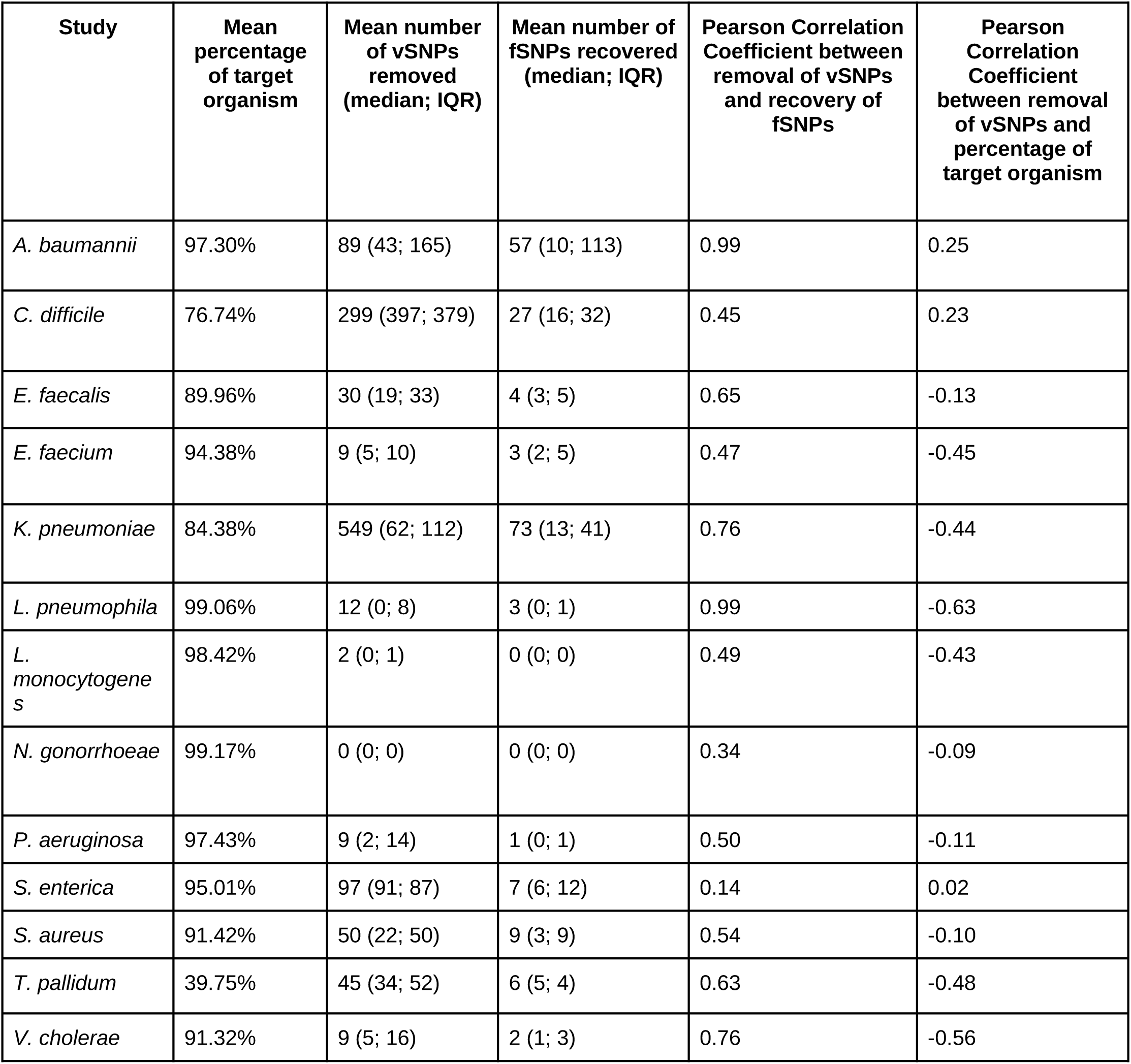
Effect of applying the taxonomic filter in the variant analysis of samples of the *bacterial dataset*.

**Figure 2:**
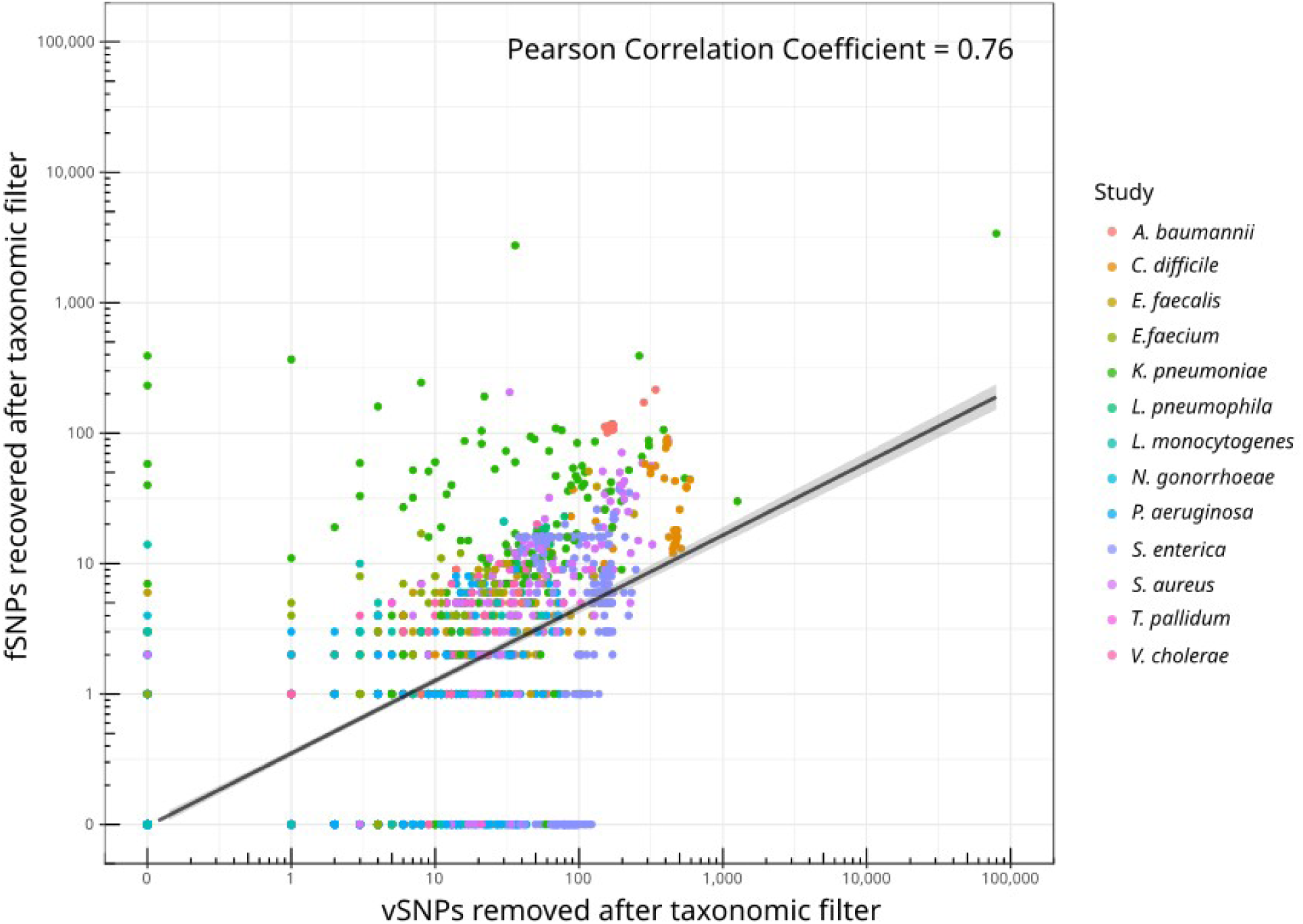
Correlation between the number of vSNPs removed and the number of fSNPs recovered after contamination removal with the taxonomic filter.

Overall, the impact of removing contaminant reads on vSNP and fSNP inference depended heavily on the species considered. For example, virtually no change was observed for *N. gonorrhoeae* samples (Table 2, Figure 3) while a mean number of 549 vSNPs were removed and 73 fSNPs recovered for *K. pneumoniae* samples. In many WGS applications genetic variants are not analyzed on a sample basis but across the entire dataset. We therefore evaluated the impact of contaminant reads on polymorphic positions called across datasets. On average, the total number of polymorphic positions was reduced by 1.51% for fSNPs (range 0% - 6%) and 8.67% for vSNPs range (0% - 41%) (Figure 3, Supplementary Table 2).

**Figure 3:**
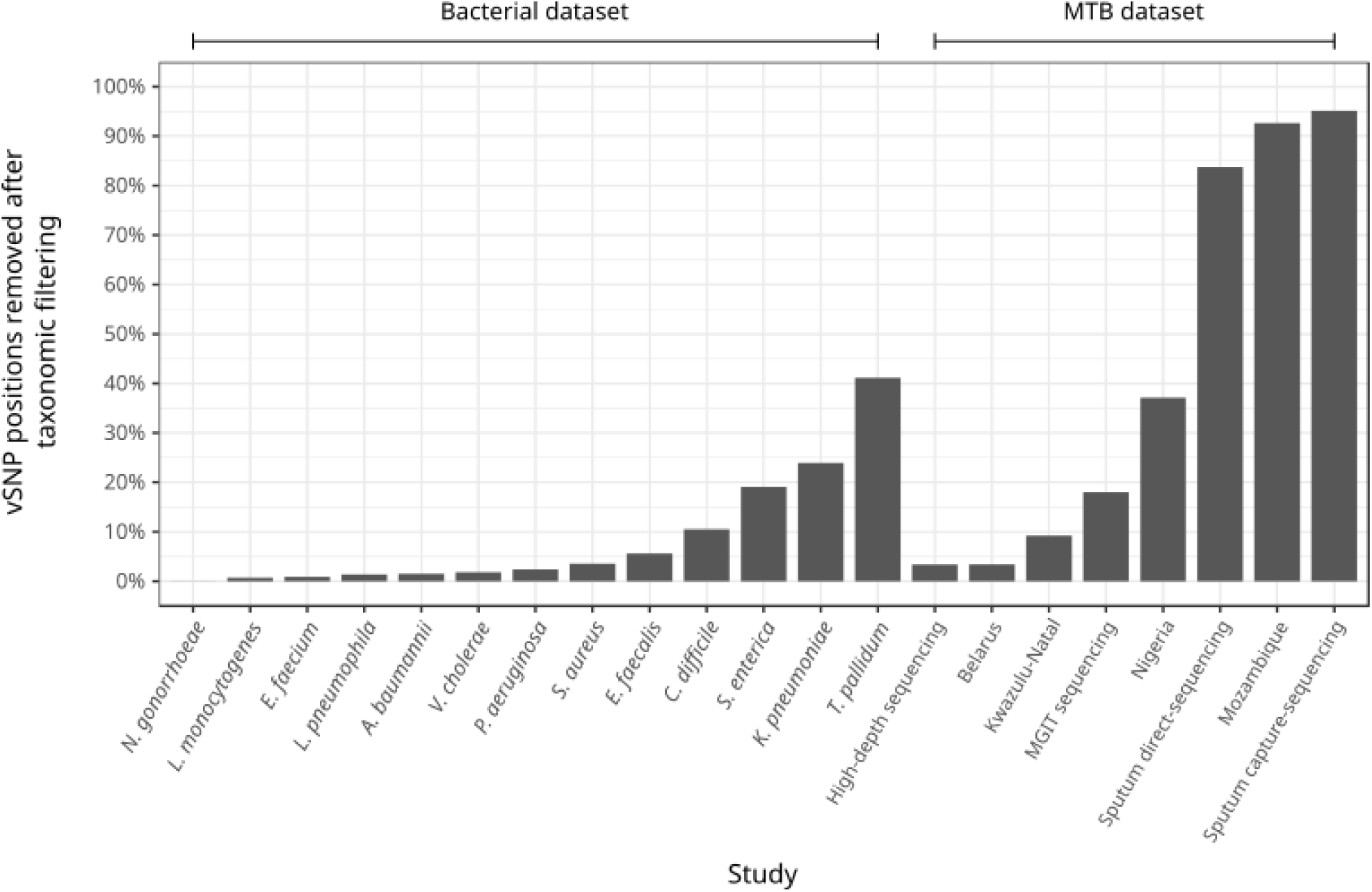
Fraction of polymorphic positions with vSNPs removed after applying the taxonomic filter for each one of the studies analyzed.

Unexpectedly, we also observed a small proportion of fSNPs to be systematically removed by the taxonomic filter (median=0.2% of fSNPs, ranging from 0% to 5.6% between studies; Supplementary Table 3). Those positions can be considered false negatives introduced by the pipeline, including inconsistencies of the mapping software, and the inability of Kraken to classify a small proportion of reads disregarding their similarity to the reference genome (further discussed in Supplementary Results 1 and Supplementary Figure 1). When inspecting a fraction of the removed fSNPs, we observed that most of them were across low coverage regions. Removing few reads in those regions makes the position fall below the required thresholds to a call a fSNP in the filtered sample.

Thus, our results not only show that contaminants have a major impact on variant analysis, but also that dealing with such contaminants will require different contamination-control strategies and specific implementations for each organism to reach an acceptable trade-off between false positives and negatives.

### Implementation of a contamination-aware analysis pipeline: *Mycobacterium tuberculosis* as a test case

Given the results observed in the *bacterial dataset*, we implemented two contamination-control approaches on top of a specific analysis pipeline for *M. tuberculosis* WGS data (see Methods): a taxonomic filter at species level (*Mycobacterium tuberculosis* complex) and a similarity filter that removes read mappings with identity and length lower than 97% and 40 bp respectively. We tested both approaches using simulated and real sequencing runs. In first instance, we used *in-silico* simulated experiments to evaluate how non-MTB reads are mapped to the MTB reference genome and quantify the false positive and negative SNPs that arise as a consequence. We mapped simulated sequencings of 45 organisms to the MTB reference genome, including oral and respiratory microbiota, clinically common non-tuberculous mycobacteria, and human reads. As expected, conserved genes like the 16S, *rpoB*, or the tRNAs, constitute hotspots where contaminant sequences are frequently aligned to. However, non-MTB alignments are not only produced in these regions but across the reference genome (Figure 4a). This is dependent on the phylogenetic relationship of the contaminant organism to the one being studied. Non-tuberculous mycobacteria represent the best example of this, as their read mappings can produce high sequencing depths along the MTB reference genome. Remarkably, human reads, which are frequently the main concern in clinical samples, did not produce alignments at all.

**Figure 4:**
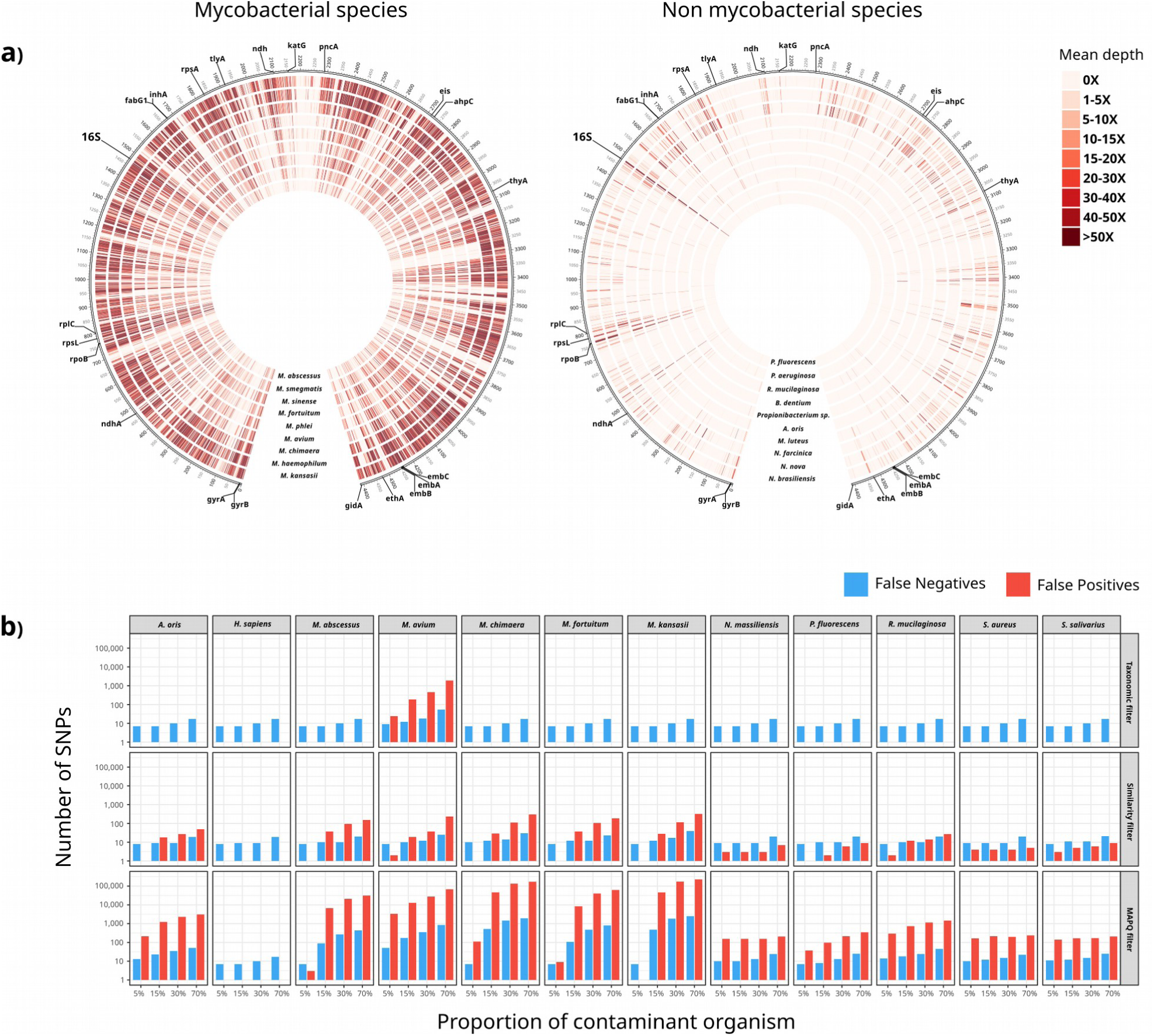
Mapping of non-MTBC reads across the MTBC reference genome impacts variant calling. **a)** Mean sequencing depth obtained along the MTBC reference genome across 1000-bp windows when mapping 1,500,000 simulated reads of non-tuberculous mycobacteria species and organisms other than mycobacteria (OTM). For OTM, the 10 organisms that produced higher sequencing depths are shown. **b)** Number of false positive vSNPs and false negative fSNPs (note logscale) in samples in-silico contaminated with different proportions of non-MTB organisms when following three different analysis pipelines (taxonomic filter, similarity filter and a standard pipeline including a mapping quality filter (MAPQ 60))Next, we evaluated the performance of the taxonomic filter and the similarity filter

using *in-silico* contaminated samples. Whereas both approaches reduced the number of non-MTB mappings, the taxonomic filter showed the best performance, eliminating all non-MTB alignments with the only exception of a proportion of *M. avium* reads. Accordingly, the number of false positive vSNPs due to contaminants was reduced with both methods, but in the case of the taxonomic filter almost all erroneous SNP calls were eliminated (Figure 4b). Only contaminations with *M. avium*, a closely related bacteria, compromised its performance. Nonetheless, the errors observed were notably lower than when only using a mapping quality threshold (60 in this work). For example, when a 5% of *M. avium* was present, the 3,325 false positive vSNPs and 51 false negative fSNPs identified were reduced to 24 and 9 respectively after applying the taxonomic filter. The few false negative fSNPs observed in Figure 4b that are systematic between all methods, were due to some positions next to hard-to-map regions that do not pass the coverage cutoffs required to call a fSNP in contaminated samples.

Remarkably, even slight contaminations (5% in this simulation) can introduce a large number of false positive vSNPs. As expected, the erroneous calls produced by such small contaminations fall mainly in conserved regions. However, in agreement with the results shown in Figure 4a, spurious SNPs can be called across the genome (Supplementary Figure 2). Importantly, it is precisely because many of the contaminant alignments are produced in conserved genes that we predicted false antibiotic resistances, including well-known mutations to first line drugs in MTB treatment (Supplementary Table 4).

We also evaluated whether these filters systematically remove sequencing reads from particular genomic regions leading to biases produced by the methodology itself. To do so, we analyzed the mean sequencing depth obtained across the genome, before and after applying the filters, for all the samples of the *MTB dataset* that have less than 1% of contamination (984 samples; 78% of the samples analyzed). Importantly, we observed the taxonomic filter to systematically remove sequencing reads coming from the 16S gene due to the inability of Kraken to classify many reads coming from this gene up to the level of species. However, for the rest of the genome it showed an excellent performance, with virtually no differences in depth, even for conserved regions like the *rpoB* gene (Supplementary Table 5). On the contrary, the similarity filter produced a systematic decrease in depth across the genome. In the 97% of the genome the sequencing depth was reduced more than 1X, with several regions showing larger decreases (Supplementary Table 6).

### Impact of contaminations in clinical WGS samples of *Mycobacterium tuberculosis*

After evaluating the performance of the taxonomic and similarity filters, we used them to remove contaminants in a dataset comprising 1,553 MTB WGS samples from eight different studies. As done for the *bacterial dataset*, we only analyzed samples with at least 50% of reads classified as *Mycobacterium tuberculosis* complex and 40x of median sequencing depth (20X for direct sequencing from clinical specimens) to discard heavily contaminated sequencings (1,267 samples, 81.6% of the *MTB dataset*)

Given that the taxonomic filter showed to be extremely conservative with all genomic positions except the 16S gene, we discarded from the following analyses any SNP called in this region (*rrs, rrl, rrf)*. Therefore, the differences observed in variant analysis when applying this filter can be attributed to noise introduced by contaminations. In accordance, we expected no differences in variant calling in samples not affected by contaminants. When analyzing real WGS MTB samples with the taxonomic filter we observed no variant change for 788 samples (62% of the samples analyzed). Importantly, this agreement was true for samples with low-level contaminations (less than 1%) but also for samples with higher number of contaminant reads (up to 31%). Overall, the number of SNPs either removed or recovered after applying the taxonomic filter were independent of the level of contamination of a sample (Pearson Correlation Coefficient = 0.03). Altogether, these results strongly support that the changes observed in variant analysis after applying the taxonomic filter can be attributed to noise introduced by contaminants rather than a methodological bias. On the contrary, the similarity filter always remove variant positions even for the 984 samples with 99% of MTB. This is in agreement with the higher rate of false negatives observed in the *in-silico* experiments.

Contaminant read mappings introduce new variants that alter the allele frequencies. After applying the taxonomic filter, we observed a mean change of 42% allele frequency (median=41%; IQR=36%). As shown in Figure 5, the main consequence of these alterations is the introduction of many false positive vSNPs, even for samples with contaminations as low as 5%. However, altering allele frequencies can also lead to call false negative vSNPs, and false positive and negative fSNPs. Among the 38% of samples for which at least one change was observed, the taxonomic filter removed on average 761.7 vSNPs (median=18) and 4 fSNPs (median=1), and recovered 1.7 vSNPs (median=1) and 5.9 fSNPs (median=2). On average, the total number of polymorphic positions within each study was reduced by 0.4% for fSNPs (range 0% - 2%) and 43% for vSNPs range (3% - 95%) (Figure 3, Supplementary Table 2). Applying the similarity filter removed on average 129.1 vSNPs (median=20) and 6.1 fSNPs (median=5) and recovered 2.6 vSNPs (median=2) and 2.3 fSNPs (median=2).

**Figure 5:**
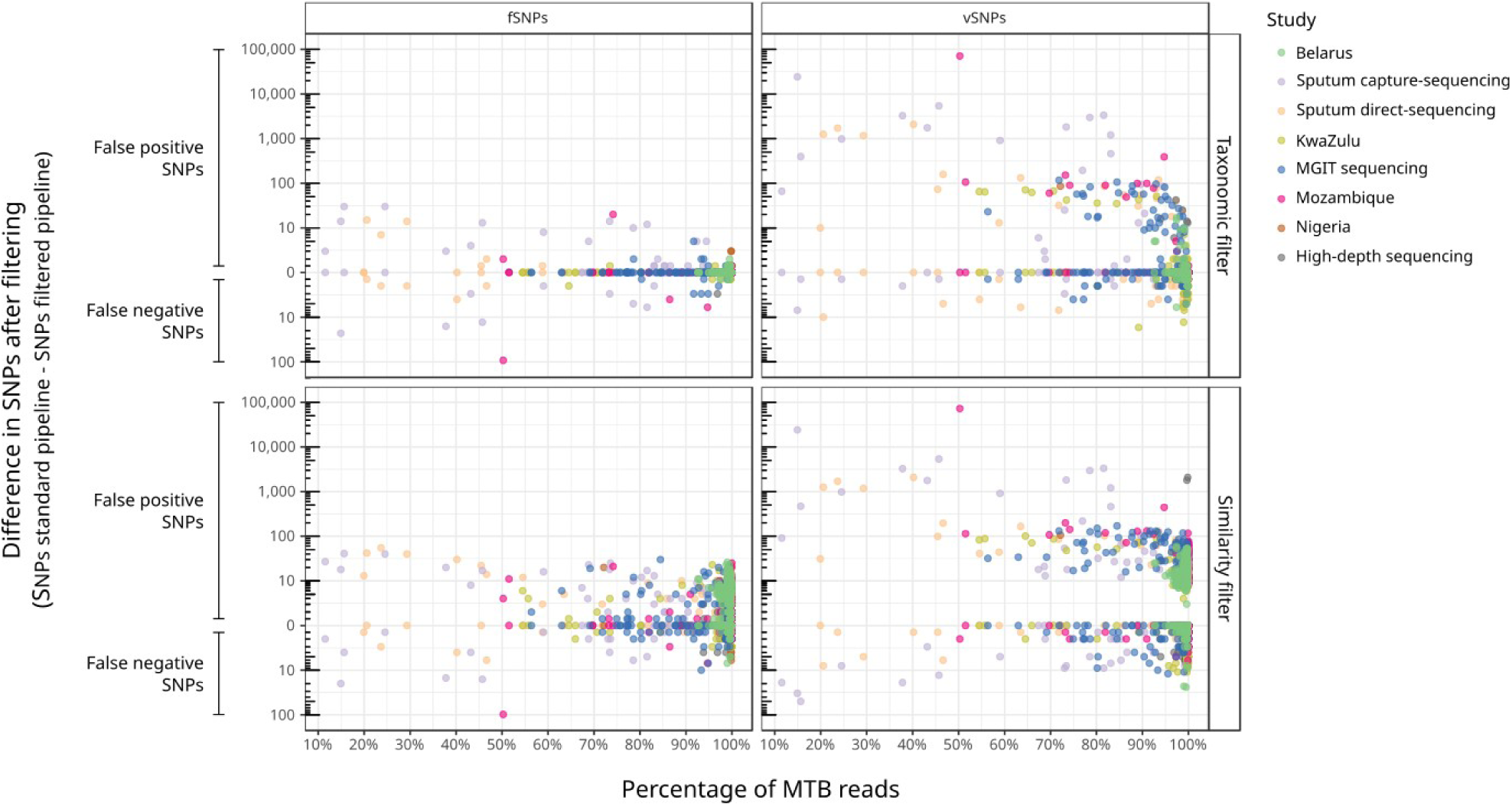
Differences in SNP calling in samples of the MTB dataset between a standard pipeline and the two contamination-control methodologies tested. Rulers at the left of the graphic highlight the false positive and negative SNPs attributable to contamination according to each filtering methodology.

Sequencing directly from clinical specimens is subject to greater alterations in variant analysis (Figure 5) since this strategy usually yields highly contaminated samples and limited sequencing depth. In these cases, the SNP frequencies are more sensitive to contaminant reads since only few reads can be responsible for a shift in the frequencies that make a position to fall below or above the required thresholds to call a variant (Supplementary Figure 1). However, a high sequencing depth does not guarantee an analysis safe of errors either. This effect can be observed in the High-depth sequencing study, a work based on low-frequency variant analysis from samples with more than 1000X sequencing depth. In this study, 7 samples out of 63 showed changes in the SNP analysis after applying the taxonomic filter. On average, 16.9 false positive vSNPs were removed (ranging from 2 to 42 vSNPs) and for one sample 3 false negative fSNPs and 2 vSNPs were recovered. Remarkably, no strong contamination was detected for these samples (with MTB ranging from 96.86% to 99.84%). For instance, in a sample with as much as 99.84% of MTB, the taxonomic filter removed 13 false positive vSNPs in 12 different genes across the genome.

## Discussion

In this work we analyze more than 4,000 WGS samples from 14 different pathogenic bacterial species to evaluate the impact of contaminations in WGS studies. We demonstrate that contaminant reads suppose a great pitfall since they are unexpectedly frequent and can have a large impact in variant analysis, which is the foundation of many genomic analyses. As expected, contaminations are a main issue when sequencing DNA that has not been extracted from pure cultures or single colonies, like in the case of clinical specimens. However, we show that experiments sequencing from pure cultures are not necessarily free of contamination, and that using standard mapping quality parameters are not enough to deal with contaminant reads. Therefore, bioinformatic pipelines assuming that all the reads successfully mapped are from the target organism might lead to a biased variant analysis.

We show that the errors introduced by contaminations are very variable among different studies, (Table 2; Figure 3; Figure 5), which differ not only on the organism being sequenced but also on the sampling source and laboratory protocols. For example, in the *T. pallidum* study, where samples are heavily contaminated, very little differences are observed in the variant analysis. This stems from the fact that most of the contamination in this study comes from human reads, unlikely to map to the *T. pallidum* genome. On the contrary, for the *L. pneumophila* dataset, a sample with 96.27% of *Legionella*, had 79 vSNPs and 5 fSNPs removed, and 17 fSNPs recovered after filtering a 3% of unclassified reads. According to the NCBI blast, a fraction of those reads was from *Legionella spiritensis*. The downstream relevance, however, is not directly proportional to the absolute number of erroneous SNPs and frequencies, but to what that errors mean for each organism. For example, for organisms with low genetic diversities, like in the case of MTB, a change in few fSNPs can have major implications in epidemiology studies since transmission cutoffs vary between 5 to 12 fSNPs(*19*). This is also true when predicting drug-resistance, particularly considering that many drug-resistance associated genes are conserved among bacteria and hence more prone to recruit contaminant mappings. Likewise, the higher impact observed for the vSNPs, both in terms of absolute numbers and frequencies, can have large implications in those applications based on the analysis of the allele frequency spectrum, for example when studying complex traits in bacterial populations.

A main limitation of this study is that we used the same bioinformatic pipeline to analyze WGS from organisms that are genetically different. This might have led us to either over or underestimate the effects of contaminations for some organisms, where species-specific filters might be needed for a more accurate analysis. For MTB, for instance, repetitive and mobile elements (accounting for ∼10% of the genome) are typically removed from the analysis. However, we think that the problem is mitigated by the fact that the analysis on the contaminants in the *bacterial dataset* is limited to reads from genera other than the target organism and, therefore, our estimation of the impact of contaminations among these species is likely to be underestimated. This is particularly true considering that the closer the contaminant organism is to the one under study, the larger the errors produced in variant analysis. Implementing accurate contamination-aware pipelines for different organisms will require specific analysis workflows that must be comprehensively evaluated to ensure the reliability of the results.

We performed such comprehensive evaluation for WGS analysis of MTB, for which we benchmarked two methods to remove contaminant mappings using both real and simulated data. Our analysis shows more accurate results using a taxonomic filter as compared to the similarity filter, what is probably true for any other organism with representative genomes in the databases and moderate genetic diversities. The analyses for MTB reveal a large number of variants introduced by contaminants with downstream consequences when calling vSNPs and fSNPs as well as the wild type. Remarkably, we show that contaminations can introduce substantial errors in samples that could be considered “pure” or with high sequencing depths, implying that contamination-aware pipelines will be needed in any circumstance.

The robustness and high-accuracy of the Kraken-based taxonomic filter for MTB had a cost of a systematic decrease of coverage in the 16S gene, what is not relevant for some applications (phylogeny, epidemiology) but is relevant for detecting resistance to some aminoglycosides. Kraken provided our implementation the necessary balance between speed and accuracy to analyze thousands of samples. However, multiple strategies and software can be used to develop taxonomic or similarity filters, depending on the necessities of each research group and application.

Contaminations suppose a usually neglected pitfall in WGS studies that can introduce large biases in variant analysis. Our results at the variant level parallels those observed in genomic repositories (*16, 18*). Therefore, we call for the use of validated contamination-aware pipelines in any bacterial WGS study. These analyses pipelines should be capable of, at least, report the contaminated samples and their contaminants to be later interpreted by the researcher. Ideally, they should be able to produce accurate results regardless of the extent of contamination of a sample. Pipelines capable of accurately analyze contaminated WGS data will soon become essential, since the improvement of laboratory protocols allows the sequencing of an increasing number of bacterial species directly from clinical specimens(*20, 21*). In this work, we provide a highly accurate contamination-aware pipeline for MTB WGS analysis that will be extremely helpful in the upcoming studies and clinical applications sequencing MTB directly from respiratory samples.

## Material and Methods

### Datasets analyzed from bacterial WGS studies

In order to detect contamination through different studies and evaluate its impact in bacterial WGS experiments, we analyzed WGS runs from 20 different studies. We considered studies that have been published recently and for which Illumina sequencing reads were already available for downloading. The datasets comprised 8 MTB studies and 12 studies of other 13 relevant pathogenic species. Nineteen of these datasets were publicly available beforehand (*22*–*40*). To include a dataset generated in our laboratory, we sequenced 138 MTB samples from Mozambique in the Illumina MiSeq platform (Supplementary Methods 1). A total of 4,194 Illumina runs were analyzed, comprising 1,553 MTB samples (*MTB dataset)* and 2,641 samples from the rest of organisms (*bacterial dataset)* (Table 1).

### Contamination assessment using Kraken

In order to assess contamination in each dataset, sequencing reads were taxonomically classified using Kraken(*41*) with a custom database comprising all sequences of bacteria, archaea, virus, protozoa, plasmids and fungi in RefSeq (release 78), plus the human genome (GRCh38, Ensembl release 81). Kraken classifications and kraken database setup were performed with default parameters.

### Analysis pipeline

To analyze WGS data we used a general analysis pipeline for read mapping and variant calling. In summary, reads were trimmed and filtered to remove low-quality sequences and then mapped to the reference genome of each organism using bwa mem (*42*). We used as reference genomes those used by the authors in their respective manuscripts when specified and otherwise the representative genome of RefSeq (Supplementary Table 7). For MTB samples we used the genome of the inferred most recent common ancestor of the *Mycobacterium tuberculosis* complex. Alignments with mapping qualities (MAPQ) below 60 were removed. Variants were then called and filtered using two different set of parameters to call fixed SNPs (fSNPs) and variable SNPs (vSNPs). The cutoffs to call fSNPs were minimum depth of 20 reads, with the variant observed in at least 20 reads, average base quality of 25, *p-value* cutoff of 0.01, observed in both strands and minimum frequency of 90%. The cutoffs to call vSNPs were minimum depth of 10 reads, with the variant observed in at least 6 reads, average base quality of 25, *p-value* cutoff 0.01, observed in both strands and minimum frequency of 10%. We also removed SNPs near indels in a window of 4 bp. For MTB samples, we used an additional annotation filter to remove SNPs in repetitive and mobile regions. Additionally, to call fSNPs, we used a density filter removing SNPs within high-density regions (allowing a maximum of 3 SNPs in 10bp windows). This filter is commonly used in MTB WGS data since it is not expected to observe many contiguous variants given the extremely low genetic diversity of this species.

We compared this general analysis pipeline with two approaches for contamination removal. The one referred as taxonomic filter consisted in the removal of contaminant reads after the trimming step, prior to mapping. For MTB samples, we removed those reads classified by Kraken as any species other than *Mycobacterium tuberculosis* complex. In the case of organisms other than MTB, to be conservative, we removed the reads classified as any organism other than the target at the level of genus, keeping also those sequences classified as phages of that organisms. For MTB samples, we also tested another method consisting in a custom *similarity filter* to eliminate low-quality alignments consisting in the removal of alignments with length and identity below 40 bp and 97% respectively.

Importantly, in this work we only considered for analysis those samples where the errors introduced by contaminations would not be easily detected with standard pipelines. Therefore, extreme biases introduced by highly contaminated sequencings are not reported in this work. To do so, we only evaluated the impact of contaminations in variant calling for samples with more than the 50% of the target organism and with a median depth of at least 40X. In the case of studies performing WGS directly in clinical samples (sputum-capture sequencing, sputum-direct sequencing and *Treponema* studies) we analyzed those samples that had at least 20X of median coverage, since in this type of sequencing is expected to sequence samples with lower coverages and high proportions of non-target reads.

### Generation of simulated datasets

We used the reference genome of each organism to generate simulated sequencing samples using ART(*43*). We generated paired-end sequencings of 250 and 100 bp using the errors profiles of Illumina MiSeq (--ss MSv3) and Illumina HiSeq (-ss HS20) platforms respectively. This allowed us to estimate the proportion of reads that cannot be classified by Kraken up to level of genus and species for each organism. The same approach was used to generate sequencing runs of different bacterial contaminants commonly observed in MTB WGS samples (see below). The command line used to generate the sample was:

~~~
art_illumina -ss [MSv3 | HS20] --id {} --rcount 2000000 –in {}.genomic.fna -l 250 --mflen 800 --out simulated_reads/{}. --paired -- minQ 25 -s 300
~~~

### Evaluation of the impact of contaminations and methodology validation

We generated sequencings for the MTB reference genome, the human genome (GRCh38, Ensembl release 81) and 44 different non-MTB bacterial species (Supplementary Table 8). This allowed us to perform two kind of experiments (mapping of non-MTB reads to the MTB reference genome and analysis of mock contaminated samples) as explained further below.

In order to inspect which regions of the reference genome are susceptible of recruiting non-MTB reads, we mapped the simulated reads and then measured the mean sequencing depth across the genome in 1000 bp windows. To assess whether false positive SNPs and drug resistance predictions are produced by these non-MTB mappings, we generated mock contaminated samples by mixing sequencing reads of the reference genome with different proportions (5%, 15%, 30% and 70%) of other organisms corresponding to 12 common contaminants identified in the *MTB dataset*. Therefore, any SNP identified when analyzing these samples would be false positive SNPs imputable to contaminations.

In addition, we mapped these mock samples to a modified version of the reference genome where we introduced random mutations each 100bp, and all the drug resistance conferring mutations described as “high confidence” in the PhyResSE catalog(*44*). Therefore, any of the introduced SNPs that were undetected when analyzing these samples, would be false negative SNPs attributable to contamination.

## Supplementary Materials

Supplementary methods 1. Whole genome sequencing of MTB samples from Mozambique.

Supplementary Results 1. Limitations of the Kraken-based taxonomic filter.

Supplementary Table 1. Evaluation of the performance of Kraken classifying reads at genus and species level for the reference genomes and among all samples of the studies analyzed.

Supplementary Table 2. Difference in the number of variant positions within a dataset between the basic and the taxonomic-filtered pipeline.

Supplementary Table 3. Proportion of fSNPs removed per sample in the *bacterial dataset*.

Supplementary Table 4. Evaluation of false drug resistance predictions in mock contaminated samples.

Supplementary Table 5. Genomic regions (1,000 bp windows) with a coverage decrease greater than 1X after taxonomic filtering in 984 samples of the *MTB dataset* with more than 99% of reads classified as MTB.

Supplementary Table 6. Top ten genomic regions (1,000 bp windows) with greater coverage decrease after applying the similarity filter in 984 samples of the *MTB dataset* with more than 99% of reads classified as MTB.

Supplementary Table 7. Reference genomes of the *bacterial dataset*.

Supplementary Table 8. Non-MTB species included in the simulated sequencings to evaluate the impact of contaminations in MTB WGS samples.

Supplementary Figure 1. Effects of contaminations and taxonomic filtering in variant calling.

Supplementary Figure 2. Contaminations can lead to incorrect calls across the *M. tuberculosis* genome.

## Acknowledgments

This work was funded by projects of the European Research Council ERC) (638553-TB-ACCELERATE) and Ministerio de Economía y Competitividad (Spanish Government) research grant SAF2016-77346-R (to IC), and BES-2014-071066 (to GAG).

## Author contributions

IC and GAG designed the study, analyzed the data and wrote the manuscript. SB and AGB provided the MTB samples from Mozambique.

## Competing interests

The authors declare no conflicts of interest in this article.

## Data and materials availability

Whole genome sequencing data from Mozambique isolates generated in our laboratory is available at the European Nucleotide Archive under the accession PRJEB27421. The inferred most recent common ancestor genome of the *Mycobacterium tuberculosis* complex is available at https://gitlab.com/tbgenomicsunit/Publications_resources/blob/master/MTB_ancestor.fas

## Notes

#### Summary of Updates

This version is a major update to the first manuscript in which we analyzed 1,500 samples of *M. tuberculosis*. We have now extended our analysis to other 13 pathogenic bacterial species and reanalyzed all the samples with additional stringent filters.

https://gitlab.com/tbgenomicsunit/Publications_resources/blob/master/MTB_ancestor.fas

## References and Notes

1. P. R. McAdam, E. J. Richardson, J. Ross Fitzgerald, High-throughput sequencing for the study of bacterial pathogen biology. Curr. Opin. Microbiol. 19, 106–113 (2014).

2. X. Didelot, R. Bowden, D. J. Wilson, T. E. A. Peto, D. W. Crook, Transforming clinical microbiology with bacterial genome sequencing. Nat. Rev. Genet. 13, 601–612 (2012).

3. D. J. Roach et al., Correction: A Year of Infection in the Intensive Care Unit: Prospective Whole Genome Sequencing of Bacterial Clinical Isolates Reveals Cryptic Transmissions and Novel Microbiota. PLoS Genet. 13, e1006724 (2017).

4. A. C. Brown, M. T. Christiansen, Whole-Genome Enrichment Using RNA Probes and Sequencing of Chlamydia trachomatis Directly from Clinical Samples. Methods Mol. Biol. 1616, 1–22 (2017).

5. D. J. SenGupta et al., Whole-genome sequencing for high-resolution investigation of methicillin-resistant Staphylococcus aureus epidemiology and genome plasticity. J. Clin. Microbiol. 52, 2787–2796 (2014).

6. J. A. Lees et al., Evaluation of phylogenetic reconstruction methods using bacterial whole genomes: a simulation based study. Wellcome Open Res. 3, 33 (2018).

7. S. D. Bentley, J. Parkhill, Genomic perspectives on the evolution and spread of bacterial pathogens. Proc. Biol. Sci. 282, 20150488 (2015).

8. D. Falush, Bacterial genomics: Microbial GWAS coming of age. Nat Microbiol. 1, 16059 (2016).

9. R. E. Lenski, Experimental evolution and the dynamics of adaptation and genome evolution in microbial populations. ISME J. 11, 2181 (2017).

10. F. Campbell, C. Strang, N. Ferguson, A. Cori, T. Jombart, When are pathogen genome sequences informative of transmission events? PLoS Pathog. 14, e1006885 (2018).

11. F. R. Fields, S. W. Lee, M. J. McConnell, Using Bacterial Genomes and Essential Genes for the Development of New Antibiotics. Biochem. Pharmacol. 134, 74 (2017).

12. X. Didelot, A. S. Walker, T. E. Peto, D. W. Crook, D. J. Wilson, Within-host evolution of bacterial pathogens. Nat. Rev. Microbiol. 14, 150–162 (2016).

13. N. D. Olson et al., Best practices for evaluating single nucleotide variant calling methods for microbial genomics. Front. Genet. 6, 235 (2015).

14. C. G. Wilson, R. W. Nowell, T. G. Barraclough, Cross-Contamination Explains “Inter and Intraspecific Horizontal Genetic Transfers” between Asexual Bdelloid Rotifers. Curr. Biol. 28, 2436–2444.e14 (2018).

15. M. Ballenghien, N. Faivre, N. Galtier, Patterns of cross-contamination in a multispecies population genomic project: detection, quantification, impact, and solutions. BMC Biology. 15(2017), doi: 10.1186/s12915-017-0366-6.

16. J. Lu, S. L. Salzberg, Removing contaminants from databases of draft genomes. PLoS Comput. Biol. 14, e1006277 (2018).

17. S. Merchant, D. E. Wood, S. L. Salzberg, Unexpected cross-species contamination in genome sequencing projects. PeerJ. 2, e675 (2014).

18. F. P. Breitwieser, M. Pertea, A. Zimin, S. L. Salzberg, Human contamination in bacterial genomes has created thousands of spurious proteins. Genome Research (2019), p. gr.245373.118.

19. C. J. Meehan et al., The relationship between transmission time and clustering methods in Mycobacterium tuberculosis epidemiology. EBioMedicine. 37, 410–416 (2018).

20. R. M. Doyle et al., Direct Whole-Genome Sequencing of Sputum Accurately Identifies Drug-Resistant Mycobacterium tuberculosis Faster than MGIT Culture Sequencing. J. Clin. Microbiol. 56(2018), doi: 10.1128/JCM.00666-18.

21. N. L. Bachmann et al., Culture-independent genome sequencing of clinical samples reveals an unexpected heterogeneity of infections by Chlamydia pecorum. J. Clin. Microbiol. 53, 1573–1581 (2015).

22. A. C. Brown et al., Rapid Whole-Genome Sequencing of Mycobacterium tuberculosis Isolates Directly from Clinical Samples. J. Clin. Microbiol. 53, 2230–2237 (2015).

23. A. A. Votintseva et al., Same-day diagnostic and surveillance data for tuberculosis via whole genome sequencing of direct respiratory samples (2016), doi: 10.1101/094789.

24. L. J. Pankhurst et al., Rapid, comprehensive, and affordable mycobacterial diagnosis with whole-genome sequencing: a prospective study.. 4, 49–58 (2016).

25. K. A. Cohen et al., Evolution of Extensively Drug-Resistant Tuberculosis over Four Decades: Whole Genome Sequencing and Dating Analysis of Mycobacterium tuberculosis Isolates from KwaZulu-Natal. PLoS Med. 12, e1001880 (2015).

26. K. R. Wollenberg et al., Whole-Genome Sequencing of Mycobacterium tuberculosis Provides Insight into the Evolution and Genetic Composition of Drug-Resistant Tuberculosis in Belarus. J. Clin. Microbiol. 55, 457–469 (2017).

27. M. Senghore et al., Whole-genome sequencing illuminates the evolution and spread of multidrugresistant tuberculosis in Southwest Nigeria. PLoS One. 12, e0184510 (2017).

28. A. Trauner et al., The within-host population dynamics of Mycobacterium tuberculosis vary with treatment efficacy. Genome Biol. 18, 71 (2017).

29. S. Willems et al., Whole-Genome Sequencing Elucidates Epidemiology of Nosocomial Clusters of Acinetobacter baumannii. J. Clin. Microbiol. 54, 2391–2394 (2016).

30. N. E. Stone et al., More than 50% of Clostridium difficile Isolates from Pet Dogs in Flagstaff, USA, Carry Toxigenic Genotypes. PLoS One. 11, e0164504 (2016).

31. G. H. Tyson, J. L. Sabo, C. Rice-Trujillo, J. Hernandez, P. F. McDermott, Whole-genome sequencing based characterization of antimicrobial resistance in Enterococcus. Pathog. Dis. 76(2018), doi: 10.1093/femspd/fty018.

32. K. E. Holt et al., Genomic analysis of diversity, population structure, virulence, and antimicrobial resistance in Klebsiella pneumoniae, an urgent threat to public health. Proc. Natl. Acad. Sci. U. S. A. 112, E3574–81 (2015).

33. V. J. Timms et al., Genome Sequencing Links Persistent Outbreak of Legionellosis in Sydney (New South Wales, Australia) to an Emerging Clone of Legionella pneumophila Sequence Type 211. Appl. Environ. Microbiol. 84(2017), doi: 10.1128/aem.02020-17.

34. S. Halbedel et al., Whole-Genome Sequencing of Recent Listeria monocytogenes Isolates from Germany Reveals Population Structure and Disease Clusters. J. Clin. Microbiol. 56(2018), doi: 10.1128/JCM.00119-18.

35. K. Yahara et al., Genomic surveillance of Neisseria gonorrhoeae to investigate the distribution and evolution of antimicrobial-resistance determinants and lineages. Microb Genom. 4(2018), doi: 10.1099/mgen.0.000205.

36. R. L. Marvig, L. M. Sommer, S. Molin, H. K. Johansen, Convergent evolution and adaptation of Pseudomonas aeruginosa within patients with cystic fibrosis. Nat. Genet. 47, 57–64 (2015).

37. P. Gymoese et al., Investigation of Outbreaks of Salmonella enterica Serovar Typhimurium and Its Monophasic Variants Using Whole-Genome Sequencing, Denmark. Emerg. Infect. Dis. 23, 1631–1639 (2017).

38. D. M. Aanensen et al., Whole-Genome Sequencing for Routine Pathogen Surveillance in Public Health: a Population Snapshot of Invasive Staphylococcus aureus in Europe. MBio. 7(2016), doi: 10.1128/mBio.00444-16.

39. M. Pinto et al., Genome-scale analysis of the non-cultivable Treponema pallidum reveals extensive within-patient genetic variation. Nat Microbiol. 2, 16190 (2016).

40. D. R. Greig et al., Evaluation of Whole-Genome Sequencing for Identification and Typing of Vibrio cholerae. J. Clin. Microbiol. 56(2018), doi: 10.1128/JCM.00831-18.

41. D. E. Wood, S. L. Salzberg, Kraken: ultrafast metagenomic sequence classification using exact alignments. Genome Biol. 15, R46 (2014).

42. H. Li, R. Durbin, Fast and accurate short read alignment with Burrows-Wheeler transform. Bioinformatics. 25, 1754–1760 (2009).

43. W. Huang, L. Li, J. R. Myers, G. T. Marth, ART: a next-generation sequencing read simulator. Bioinformatics. 28, 593–594 (2012).

44. S. Feuerriegel et al., PhyResSE: a Web Tool Delineating Mycobacterium tuberculosis Antibiotic Resistance and Lineage from Whole-Genome Sequencing Data. J. Clin. Microbiol. 53, 1908–1914 (2015).

