## Supplementary Material for "Contaminant DNA in bacterial sequencing experiments is a major source of false genetic variability"

### Supplementary Materials

#### Table of Contents

**Supplementary methods 1 -** Whole genome sequencing of MTB samples from Mozambique.

**Supplementary Results 1 -** Limitations of the Kraken-based taxonomic filter.

**Supplementary Table 1 -** Evaluation of the performance of Kraken classifying reads at genus and species level for the reference genomes and among all samples of the studies analyzed.

**Supplementary Table 2 -** Difference in the number of variant positions within a dataset between the basic and the taxonomic-filtered pipeline.

**Supplementary Table 3 -** Proportion of fSNPs removed per sample in the *bacterial dataset.*

**Supplementary Table 4 -** Evaluation of false drug resistance predictions in mock contaminated samples.

**Supplementary Table 5 -** Genomic regions (1,000 bp windows) with a coverage decrease greater than 1X after taxonomic filtering in 984 samples of the *MTB dataset* with more than 99% of reads classified as MTB.

**Supplementary Table 6 -** Top ten genomic regions (1,000 bp windows) with greater coverage decrease after applying the similarity filter in 984 samples of the *MTB dataset* with more than 99% of reads classified as MTB.

**Supplementary Table 7 -** Reference genomes of the *bacterial dataset*.

**Supplementary Table 8 -** Non-MTB species included in the simulated sequencings to evaluate the impact of contaminations in MTB WGS samples.

**Supplementary Figure 1 -** Effects of contaminations and taxonomic filtering in variant calling.

**Supplementary Figure 2 -** Contaminations can lead to incorrect calls across the *M. tuberculosis* genome.

**Supplementary methods 1. Whole genome sequencing of MTB samples from Mozambique.**

DNA extractions were performed in heat-inactivated samples of MTB Löwenstein-Jensen cultures with an automated DNA extraction platform (NucliSENS EasyMag; bioMérieux). Sequencing libraries were prepared with Nextera XT DNA Library Preparation Kit v3 (Illumina, San Diego, CA) following the manufacturer instructions. Whole genome sequencing was performed on the Illumina MiSeq instrument with 2X300bp paired-end reads.

**Supplementary Results 1 - Limitations of the taxonomic filter**

In the *bacterial dataset*, we observed a small proportion of fSNPs to be systematically removed after applying the taxonomic filter (Supplementary Table 3). We observed this to occur mostly in low coverage regions that could be the result of, for instance, hard-to-map regions like repetitive elements or different strains coexisting in the same sample one of which has a deletion. In such regions, eliminating only one read can lead to greater differences in frequencies making the position fall below the required thresholds to call a fSNP (Supplementary Figure 1a-c). Most of the fSNPs incorrectly removed after the taxonomic filter were caused by the inability of Kraken to classify some reads up to the level of genus. This behaviour is a known limitation of the taxonomic classifiers for conserved regions among bacteria, as we showed for the 16S in the *MTB dataset.* However, in many cases the incorrectly eliminated sequences corresponded to reads mapped with 100% identity to the reference genome and surrounded by other sequences that were classified, even at the level of species, despite having several SNPs (Supplementary Figure 1d). This is probably due to the fact that, for some reads, the k-mers in which they are decomposed, do not allow Kraken to classify beyond a given taxonomic level, disregarding the sequence diversity. However, since the immediate contiguous reads can be correctly classified, this bias is compensated by the sequencing depth.

Unexpectedly, we also observed an inconsistency of the mapping software (bwa mem) to be responsible for a small fraction of the SNPs that are either removed or recovered after applying the taxonomic filter. In these cases, we observed that the number of supporting reads at a given position differed in one read despite the fact that none of the reads mapping to that position were classified as contaminant. Surprisingly, we observed this to be the result of the exact same read, mapping to the same genomic position, with different qualities for the filtered and non-filtered fastq files. The fact that the fastq files are different (because some reads are removed in the filtered fastq), makes bwa to produce different results for a small number of reads (between 1 and 6 in our tests), that are filtered because of the mapping quality cutoff (60). We confirmed this behaviour by randomly sorting and mapping different times a set of fastq files (data not shown; tested versions 7.10, 7.12 and 7.17).

Notably, *Negativicoccus massiliensis* appeared in high proportions in many studies. This is most likely due to a bias in the databases, in which probably there is a lack of genomes from organisms that are genetically close to this species. Similarly, Kraken left a high proportion of reads unclassified in many samples. This could be mainly due to either the absence of the organism from the database, or sequences that Kraken cannot classify up to the level of genus, for instance when analyzing organisms with high genetic diversity. Indeed, when using the NCBI blastn to search a random subset of unclassified reads in the non-redundant database (nr), we observed three main patterns. Reads that either did not produce significant matches with any organism, or came from eukaryotes not present in our Kraken database; reads that produced partial alignments with many different taxa; and reads that produced good alignments, even with the target organism, but having alignment identities below 90%, what makes Kraken unable to find exact matches of 31 base pairs.

**Supplementary Table 1.** Evaluation of the performance of Kraken classifying reads at genus and species level for the reference genomes and among all samples of the studies analyzed.

| **Organism** | **Reads classified as target species (Illumina MiSeq; 250bp)** | **Reads classified as target genus (Illumina MiSeq; 250bp)** | **Reads classified as target species (Illumina HiSeq; 100bp)** | **Reads classified as target genus (Illumina HiSeq; 100bp)** | **Maximum proportion classified among all samples from each corresponding study as target species** | **Maximum proportion classified among all samples from each corresponding study as target genus** |
|---|---|---|---|---|---|---|
| *A. baumannii* | 99.07% | 99.61% | 97.98% | 99.54% | 97.37% | 99.39% |
| *C. difficile* | 99.40% | 99.40% | 98.95% | 98.95% | 98.37% | 98.37% |
| *E. faecalis* | 99.55% | 99.83% | 99.07% | 99.65% | 98.11% | 98.27% |
| *E. faecium* | 99.30% | 99.95% | 98.74% | 99.79% | 97.7% | 98.97% |
| *K. pneumoniae* | 97.86% | 98.96% | 94.20% | 98% | 94.96% | 97.07% |
| *L. pneumophila* | 99.80% | 100% | 99.61% | 99.98% | 99.80% | 99.84% |
| *L. monocytogenes* | 99.26% | 99.96% | 98.57% | 99.87% | 98.32% | 99.90% |
| *M. tuberculosis* complex | 99.98% | 100% | 99.98% | 100% | 99.99% | 100% |

| | | | | | | |
|---|---|---|---|---|---|---|
| *N. gonorrhoeae* | 99.16% | 100% | 94.96% | 99.99% | 98.72% | 99.98% |
| *P. aeruginosa* | 99.95% | 99.99% | 99.85% | 99.95% | 99.79% | 99.92% |
| *S. enterica* | 99.58% | 99.73% | 98.83% | 99.25% | 98.90% | 99.25% |
| *S. aureus* | 95.01% | 95.39% | 94.57% | 95.35% | 96.10% | 95.15% |
| *T. pallidum* | 93.54% | 100% | 72.74% | 100% | 94.75% | 94.75% |
| *V. cholerae* | 99.59% | 99.83% | 98.90% | 99.74% | 98.70% | 99.82% |

* species in the case of MTB (*Mycobacterium tuberculosis* complex)

**Supplementary Table 2. Difference in the number of variant positions within a dataset between the basic and the taxonomic-filtered pipeline.** *E. faecalis and E. faecium are analyzed by separate but belong to the same study. ** In the Mozambique study one sample contaminated with 20% of *Mycobacterium sinense* contributed with 72,354 variant positions. The difference observed for the Mozambique study when disregarding this sample was of 927 vSNP positions (20.1%).

| **Study** | **Dataset** | **Difference in fSNP positions** | **Difference in vSNP positions** |
|---|---|---|---|
| *A. baumannii* | *Bacterial dataset* | 600 (0.15%) | 129 (1.52%) |
| *C. difficile* | *Bacterial dataset* | 3,624 (5.84%) | 1,437 (10.53%) |
| *E. faecalis** | *Bacterial dataset* | 3,676 (1.94%) | 635 (5.61%) |
| *E. faecium** | *Bacterial dataset* | 492 (0.25%) | 195 (0.89%) |
| *K. pneumoniae* | *Bacterial dataset* | 6,828 (1.17%) | 68,246 (23.93%) |
| *L. pneumophila* | *Bacterial dataset* | 222 (0.12%) | 157 (1.37%) |
| *L. monocytogenes* | *Bacterial dataset* | 68 (0.03%) | 53 (0.71%) |
| *N. gonorrhoeae* | *Bacterial dataset* | 0 (0%) | 2 (0.03%) |
| *P. aeruginosa* | *Bacterial dataset* | 299 (0.09%) | 276 (2.38%) |
| *S. enterica* | *Bacterial dataset* | 813 (2.92%) | 645 (19.11%) |
| *S. aureus* | *Bacterial dataset* | 3,652 (2.14%) | 2,082 (3.57%) |
| *T. pallidum* | *Bacterial dataset* | 26 (4.47%) | 252 (41.17%) |
| *V. cholerae* | *Bacterial dataset* | 2,190 (0.55%) | 220 (1.84%) |
| Kwazulu-Natal | *MTB dataset* | 1 (0%) | 240 (9.22%) |
| Nigeria | *MTB dataset* | 3 (0.32%) | 170 (37.11%) |
| Belarus | *MTB dataset* | 0 (0%) | 71 (3.42%) |
| Mozambique | *MTB dataset* | 39 (0.32%) | 71,309 (92.65%) ** |
| High-depth sequencing | *MTB dataset* | 0 (0%) | 161 (3.40%) |

| | | | |
|---|---|---|---|
| Sputum capture-sequencing | *MTB dataset* | 124 (2.19%) | 28,452 (95.11%) |
| Sputum direct-sequencing | *MTB dataset* | 24 (0.49%) | 3,686 (83.79%) |
| MGIT sequencing | *MTB dataset* | 19 (0.11%) | 517 (18%) |

**Supplementary Table 3 -** Proportion of fSNPs removed per sample in the *bacterial dataset.*

| **Study** | **Proportion of fSNPs removed by the taxonomic filter (median)** |
|---|---|
| *A. baumannii* | 0.22% |
| *C. difficile* | 0.72% |
| *E. faecalis* | 0.52% |
| *E. faecium* | 0.16% |
| *K. pneumoniae* | 0.81% |
| *L. pneumophila* | 0.00% |
| *L. monocytogenes* | 0.03% |
| *N. gonorrhoeae* | 0.00% |
| *P. aeruginosa* | 0.00% |
| *S. enterica* | 5.58% |
| *S. aureus* | 1.89% |
| *T. pallidum* | 0.00% |
| *V. cholerae* | 0.32% |

**Supplementary table 4.** Evaluation of false drug resistance predictions in mock contaminated samples.

| **Contaminant** | **Proportion** | **Mutation** | **Gene; Drug** |
|---|---|---|---|
| *R. mucilaginosa* | Starting at 5% | 1473247 C>A (HC, new) | rrs (16S); AMK/KAN/CPR |
| | | 1473329 G>T (HC; described) | |
| *A. oris* | Starting at 15% | 761154 T>A (LC, new) | rpoB; RMP |
| | | 761155 C>G (HC, described) | |
| *M. abscessus* | Starting at 15% | 761098 G>C (LC, described) | rpoB; RMP |

| | | | |
|---|---|---|---|
| *M. avium* | Starting at 15% | 6738 C>A (HC, described) | gyrB; FQ |
| | | 6742 A>G (HC, new) | |
| | Starting at 70% | 761098 G>C (LC, described) | rpoB; RMP |
| *M. chimaera* | Starting at 15% | 6742 A>G (HC, new) | gyrB; FQ |
| | Starting at 30% | 1834852 C>G (LC, new) | rpsA; PZA |
| | | 4243245 G>C (LC, new) | embA; EMB |
| | | 4247729 G>C (HC, new) | embB; EMB |
| | | 4247730 G>C (HC, described) | |
| | Starting at 70% | 2289100 T>G (HC, new) | pncA; PZA |
| *M. fortuitum* | Starting at 15% | 761111 C>T (LC, new) | rpoB; RMP |
| | Starting at 30% | 1834852 C>G (LC, new) | rpsA; PZA |
| | Starting at 70% | 761098 G>C (LC, described) | rpoB; RMP |
| | | 1918494 T>C (LC, new) | tlyA; CPR |
| *M. kansasii* | Starting at 15% | 761098 G>C (LC, described) | rpoB; RMP |
| | | 1834852 C>G (LC, new) | rpsA; PZA |
| | Starting at 30% | 781687 | rpsL; SM |
| | Starting at 70% | 1918494 T>C (LC, new) | tlyA; CPR |

False drug-resistance predictions according to high confidence (HC) and low confidence (LC) known mutations in the PhyResSE catalog. SNPs tagged as *new* correspond to undescribed variants produced in positions known to harbour drug-resistance conferring mutations.AMK (Amikacin); CPR (Capreomycin); EMB (Ethambutol); ETH (Ethionamide); FQ (Fluoroquinolones); INH (Isoniazid); KAN (Kanamycin); LZD (Linezolid); PAS (Para-aminosalicylic acid); PZA (Pyrazinamide); RMP (Rifampicin); SM (Streptomycin).

**Supplementary Table 5** - Genomic regions (1,000 bp windows) with a coverage decrease greater than 1X after taxonomic filtering for 984 samples of the *MTB dataset* with more than 99% of reads classified as MTB.

| **Region** | **Sequencing depth difference (mean)** | **Annotation** |
|---|---|---|
| 1472000:1472999 | 113.82 | *rrs* |
| 1475000:1475999 | 20.52 | *rrl* |

| | | |
|---|---|---|
| 1473000:1473999 | 12.56 | *rrs* |
| 1476000:1476999 | 8.53 | *rrl* |
| 1474000:1474999 | 3.47 | *rrl* |
| 1471000:1471999 | 2 | *murA* |
| 3705000:3705999 | 1.66 | *sdhA* |
| 1649000:1649999 | 1.28 | Rv1461 |
| 932000:932999 | 1.13 | Intergenic |

**Supplementary Table 6** - Top ten genomic regions (1,000 bp windows) with greater coverage decrease after applying the similarity filter for 984 samples of the *MTB dataset* with more than 99% of reads classified as MTB.

| **Region** | **Sequencing depth difference (mean)** | **Annotation** |
|---|---|---|
| 2266000:2266999 | 49.45 | Rv2019,Rv2020c,Rv2021c |
| 336000:336999 | 26.73 | *PE-PGRS3,PE-PGRS4* |
| 4383000:4383999 | 23.86 | *Rv3897c,Rv3898c* |
| 2137000:2137999 | 19.74 | *Rv1887,Rv1888c* |
| 3119000:3119999 | 19.27 | *Rv2813. Intergenic region(Rv2813-Rv2814c)* |
| 3750000:3750999 | 18.89 | *Rv3347c (PPE Family protein)* |
| 467000:467999 | 18.35 | *Rv0387c, Rv0388c (PPE Family protein)* |
| 2421000:2421999 | 17.46 | Rv2159c, Rv2160c |
| 3843000:3843999 | 17.16 | Rv3426 (PPe Family protein), Rv3427c |
| 3296000:3296999 | 17.13 | Rv2946c, Rv2947c (pks15) |

**Supplementary Table 7.** Reference genomes of the *bacterial dataset*.

| Dataset | Organism | Reference |
|---|---|---|
| *Acinetobacter* | *A. baumannii* | CP_000521.1 |
| *Clostridium* | *C. difficile* | NC_009089.1 |
| *Enterococcus* | *E. faecalis* | NC_004668.1 |
| *Enterococcus* | *E. faecium* | NC_017960.1 |
| *Klebsiella* | *K. pneumoniae* | AP006725.1 |
| *Listeria* | *L. monocytogenes* | NC_003210.1 |
| *Legionella* | *L. pneumophila* | NC_002942.5 |
| *Neisseria* | *N. gonorrhoeae* | GCF_900087815.1 |
| *Pseudomonas* | *P. aeruginosa* | NC_002516.2 |
| *Staphylococcus* | *S. aureus* | NC_007795.1 |
| *Salmonella* | *S. enterica* | NC_003197.2 |
| *Treponema* | *T. pallidum* | NZ_CP003679.1 |
| *Vibrio* | *V. cholerae* | NC_002505.1 & NC_002506.1 |

**Supplementary Table 8.** Non-MTB species included in the simulated sequencings to evaluate the impact of contaminations in MTB WGS samples

| Species | RefSeq Assembly Accession_Version |
|---|---|
| *Homo sapiens* | GRCh38, Ensembl release 81 |
| *Acinetobacter baumannii* | GCF_000746645.1_ASM74664v1 |
| *Actinomyces oris* | GCF_001553935.1_ASM155393v1 |
| *Atopobium parvulum* | GCF_000024225.1_ASM2422v1_V2 |
| *Bacillus cereus* | GCF_000007825.1_ASM782v1 |
| *Bacillus thuringiensis* | GCF_000497525.1_ASM49752v2 |
| *Bifidobacterium dentium* | GCF_001042595.1_ASM104259v1 |
| *Campylobacter concisus* | GCF_003049735.1_ASM304973v1 |
| *Corynebacterium pseudodiphtheriticum* | GCF_000688415.1_ASM68841v1 |
| *Enterobacter cloacae* | GCF_000025565.1_ASM2556v1 |

| | |
|---|---|
| *Enterococcus faecalis* | GCF_000007785.1_ASM778v1 |
| *Eubacterium aggregans* | GCF_900107815.1_IMG-taxon_2642422588 |
| *Fusobacterium sp.* | GCF_900015295.1_clos_1_1 |
| *Gemella sp. (oral taxon)* | GCF_001553915.1_ASM155391v1 |
| *Granulicatella elegans* | GCF_000162475.2_Gran_ele_ATCC_700633_V2 |
| *Klebsiella pneumoniae* | GCF_000240185.1_ASM24018v2 |
| *Micrococcus luteus* | GCF_000023205.1_ASM2320v1 |
| *Mycobacterium abscessus* | GCF_000069185.1_ASM6918v1 |
| *Mycobacterium avium* | GCF_000240505.1_ASM24050v2 |
| *Mycobacterium chimaera* | GCF_900116695.1_ASM90011669v1 |
| *Mycobacterium fortuitum* | GCF_001307545.1_ASM130754v1 |
| *Mycobacterium haemophilum* | GCF_000340435.2_ASM34043v3 |
| *Mycobacterium kansasii* | GCF_000157895.3_ASM15789v2 |
| *Mycobacterium phlei* | GCF_001583415.1_ASM158341v1 |
| *Mycobacterium sinense* | GCF_001667945.1_ASM166794v1 |
| *Mycobacterium smegmatis* | GCF_000015005.1_ASM1500v1 |
| *Negativicoccus massiliensis* | GCF_900155405.1_PRJEB18760 |
| *Neisseria perflava* | GCF_002863305.1_ASM286330v1 |
| *Nocardia brasiliensis* | GCF_000250675.2_ASM25067v3 |
| *Nocardia farcinica* | GCF_001182745.1_NCTC11134 |
| *Nocardia nova* | GCF_000523235.1_ASM52323v1 |
| *Porphyromonas sp. (oral taxon)* | GCF_000292995.1_Psp279F0450v1.0 |
| *Prevotella sp. (oral taxon)* | GCF_000163055.2_ASM16305v2 |
| *Propionibacterium sp. (oral taxon)* | GCF_001717565.1_ASM171756v1 |
| *Pseudomonas aeruginosa* | GCF_000006765.1_ASM676v1 |

| | |
|---|---|
| *Pseudomonas fluorescens* | GCF_000237065.1_ASM23706v1 |
| *Rothia mucilaginosa* | GCF_000011025.1_ASM1102v1 |
| *Staphylococcus aureus* | GCF_000013425.1_ASM1342v1 |
| *Streptococcus mitis* | GCF_000027165.1_ASM2716v1 |
| *Streptococcus oralis* | GCF_002871315.1_ASM287131v1 |
| *Streptococcus parasanguinis* | GCF_000164675.2_ASM16467v2 |
| *Streptococcus pneumoniae* | GCF_000007045.1_ASM704v1 |
| *Streptococcus salivarius* | GCF_000785515.1_ASM78551v1 |
| *Veillonella sp. (oral taxon)* | GCF_000221605.1_ASM22160v1 |
| *Yersinia enterocolitica* | GCF_000009345.1_ASM934v1 |

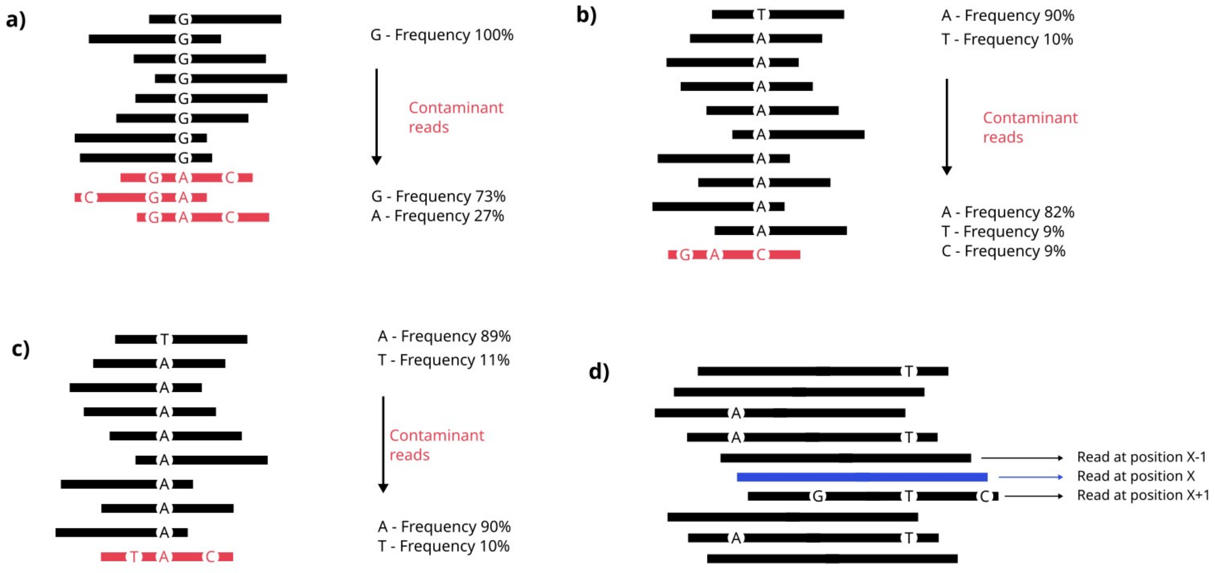


**Supplementary Figure 1. Effects of contaminations and taxonomic filtering in variant calling.** In this figure we exemplified possible effects of contaminant reads (in red) in variant calling (**a-c**) and the inability of Kraken classifying some reads (**d**). **a)** Contaminant reads make G to be called as vSNP instead of fSNP an introduce a false positive vSNP (A) **b)** Contaminant reads make A to be called as fSNP instead of a vSNP. **c)** Contaminant reads make A to be called as vSNP instead of a fSNP and make one vSNP (T) to be lost due to SNP calling cutoffs (10% frequency for vSNPs). **d)** The read coloured in blue cannot be classified by Kraken up to the level of genus. This occurs despite the fact that this read is 100% identical to the reference and that surrounding reads, with several SNPs, can be classified even to the level of species.

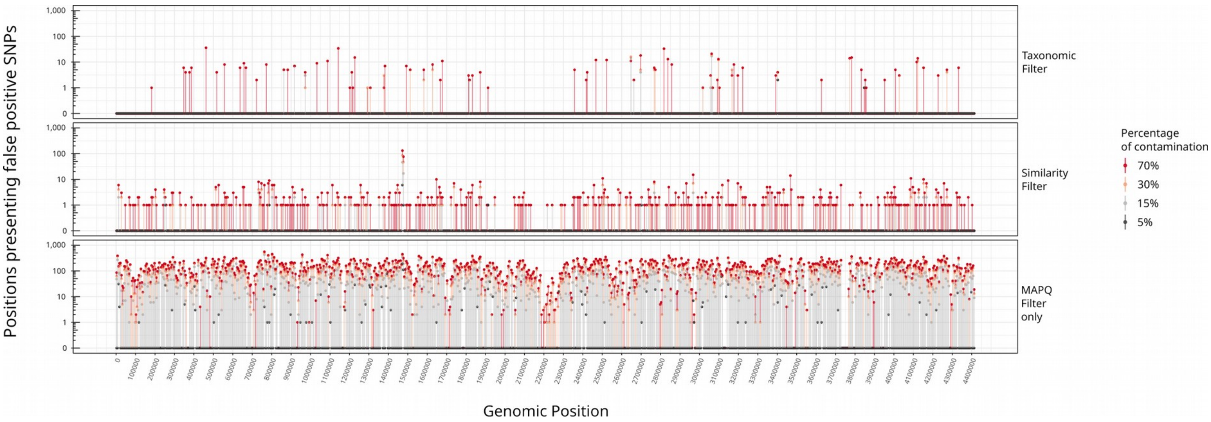


**Supplementary Figure 2. Contaminations can lead to incorrect calls across the *M. tuberculosis* genome.** Number of positions (in 1,000 bp windows) with false positive SNPs arising from 5%, 15%, 30% and 70% contaminations with different organisms in mock contaminated MTB WGS samples. Remarkably, for 5% contaminations there are some regions were false positive SNPs can be called despite the threshold of minimum frequency of 10% to call vSNPs.
